# Aberrant ciliogenesis induced by enhanced BMP signaling causes heterotopic ossification

**DOI:** 10.64898/2026.07.02.735922

**Authors:** Hiroyuki Yamaguchi, Jianbo Wang, Fangfang Yan, Jiarui Bi, Radbod Darabi, William R Lagor, Zhongming Zhao, Aris N Economides, Yuji Mishina, Yoshihiro Komatsu

## Abstract

Bone morphogenetic protein (BMP) signaling is a principal driver of heterotopic ossification (HO), yet how aberrant BMP activity structurally reprograms cellular signaling machinery to develop HO remains unclear. Here, we identify BMP signaling as a direct upstream regulator of ciliogenesis that coordinates a multi-stage, pro-osteochondrogenic signaling relay during HO. Using a conditional gain-of-function BMP mouse model (*Acvr1^Q207D/+^*), we demonstrate that enhanced BMP signaling promotes primary cilium biogenesis and axonemal elongation through canonical Smad1/5/9-dependent transcriptional activation of intraflagellar transport (IFT) *Ift20*, a core component of the IFT machinery. Rather than operating via a singular downstream cascade, these elongated cilia establish a sensitized signaling hub. Genetic disruption of ciliary Hedgehog (Hh) transduction via *Smoothened* (*Smo*) deletion reveals that ciliary Hh signaling is dispensable for initial tissue condemnation but required for the subsequent proliferative expansion and maturation of HO. Conversely, complete genetic ablation of the ciliary structure via *Ift20* deletion, or early pharmacological inhibition of ciliogenesis, significantly attenuates HO. Notably, this BMP-IFT20-cilia axis is functionally conserved within injury-responsive, PDGFRα-positive fibro-adipogenic progenitor (FAP) populations harboring the clinically authentic *Acvr1^R206H/+^* mutation responsible for fibrodysplasia ossificans progressiva (FOP) in mice. Together, these findings reveal that BMP signaling drives HO by structurally expanding the primary cilium, establishing a novel mechanism for HO development.

**Significance:** Growth factor signaling instructs cellular behavior during tissue regeneration, but how they regulate cellular organelles to induce pathological fates remains poorly understood. This study reveals that Bone Morphogenetic Protein (BMP) signaling functions as a direct architectural regulator of the primary cilium, a critical cellular antenna. We show that BMP signaling directly transactivates intraflagellar transport machinery to structurally elongate the cilium, creating a sensitized signaling hub that drives heterotopic ossification. Our findings introduce a novel BMP-driven organelle regulation mechanism and establish a targetable cellular vulnerability to mitigate ectopic bone formation.

## Introduction

Heterotopic ossification (HO) is a debilitating pathological process characterized by the *de novo* formation of extraskeletal bone within soft connective tissues (1, 2). Whether precipitated by severe mechanical trauma or driven by rare, recurrent autosomal dominant mutations within the BMP type I receptor ACVR1, such as in fibrodysplasia ossificans progressiva (FOP), HO progressively immobilizes joints, resulting in severe functional morbidity (3, 4). At the cellular level, this ectopic bone formation typically recapitulates endochondral ossification, a developmental cascade wherein a transient cartilaginous template is systematically deposited and subsequently replaced by mature mineralized bone (5–13). Following soft-tissue injury, disruption of local extracellular matrix architecture triggers a robust, localized release of osteogenic morphogens, most notably bone morphogenetic proteins (BMPs) (14–16). While the physiological roles of BMPs in skeletal patterning and development are firmly established, a fundamental mechanistic question remains unresolved: how aberrant, sustained BMP signaling instructs resident multipotent progenitors to permanently adopt a skeletogenic fate, and how secondary signaling inputs are coordinated to sustain unchecked ectopic bone growth.

Accumulating evidence indicates that the precise spatiotemporal orchestration of complex developmental signals relies heavily on the primary cilium, a solitary, microtubule-based organelle that acts as a specialized cellular antenna protruding from the vertebrate cell surface (17–19). Primary cilia serve as indispensable hubs for major morphogenetic pathways, including Hedgehog (Hh), Wnt, TGF-β, and platelet-derived growth factor (PDGF) signaling (18, 20). Consequently, decoding the mechanisms governing ciliogenesis has become a focal point in cell biology, as structural or functional ciliary defects underlie a broad spectrum of human diseases known as ciliopathies (21). Although fibroblast growth factor (FGF) signaling has been implicated in modulating ciliary length during embryonic and skeletal patterning (22–26), it remains unknown whether other major growth factor pathways actively manipulate ciliary biogenesis. Crucially, the potential intersection between osteogenic morphogens and primary cilium dynamics during pathological tissue remodeling remains completely unexplored.

Here, we demonstrate that BMP signaling acts as a direct upstream transcriptional regulator of ciliogenesis, serving to establish a multi-stage signaling relay during the pathogenesis of HO. We show that activated BMP signaling upregulates essential intraflagellar transport (IFT) machinery (17, 27), driving primary cilia assembly and structural elongation in injury-primed mesenchymal progenitor populations. By identifying BMP as a novel ciliogenic growth factor, these findings reveal a structural, cilia-mediated positive feedback loop. Mechanistically, we demonstrate that this ciliary amplification operates via a dual-phase model: while the physical presence of the ciliary axoneme is strictly epistatic for early pro-osteochondrogenic lineage commitment, its specialized downstream transducer, Hh signaling, is sequentially deployed to drive the later proliferative expansion of the HO. Ultimately, this work uncovers a unique cellular etiology for HO and establishes a new concept for how BMP pathways and organelle biogenesis converge to dictate pathological tissue identity.

## Results

### Enhanced BMP signaling drives primary cilium elongation

To investigate the transcriptional landscape initiating HO, we utilized a transgenic mouse model permitting conditional, Cre-inducible expression of a constitutively active ACVR1 variant (Acvr1^Q207D^) (28). Intramuscular injection of cardiotoxin (CTX) and adenovirus-expressing Cre recombinase (Ad-Cre) into the hindlimbs of *Acvr1^Q207D/+^* mice (Figure S1A) induced robust, localized HO by day post-injury 7 (dpi7), as confirmed by micro-computed tomography (µCT) analysis (Figure S1B, C).

Given our previous finding that genetic loss of *Acvr1* impairs node ciliogenesis during early embryogenesis (29, 30), we hypothesized that pathological activation of BMP signaling via ACVR1 may conversely augment ciliary biogenesis during HO development. To capture early molecular changes before the histological appearance of ectopic cartilage or bone, RNA sequencing (RNA-seq) was performed on hindlimb tissue lysates at dpi5. Differential expression analysis filtered through the Ciliary genes, which were defined by the SYSCILIA Gold Standard version 1 database (31), revealed the significant upregulation of 137 primary cilium-associated transcripts in Ad-Cre/CTX-treated tissues relative to Ad-GFP/CTX controls (Figure 1A). To investigate whether increased expression of ciliary genes leads to enhanced ciliogenesis, we isolated subcutaneous mesenchymal progenitor cells (SMPs) expressing PDGFRα, a surface marker expressed by the fibro-adipogenic progenitor (FAP) populations known to contribute to the HO lineage (7, 32, 33). *In vitro* treatment of *Acvr1^Q207D/+^* SMPs with Ad-Cre resulted in a significant, cell-autonomous increase in both primary cilium prevalence and total axoneme length compared with Ad-GFP controls, confirming that hyperactivated ACVR1 signaling directly stimulates ciliary elongation (Figure 1B, C).

**Figure 1.**
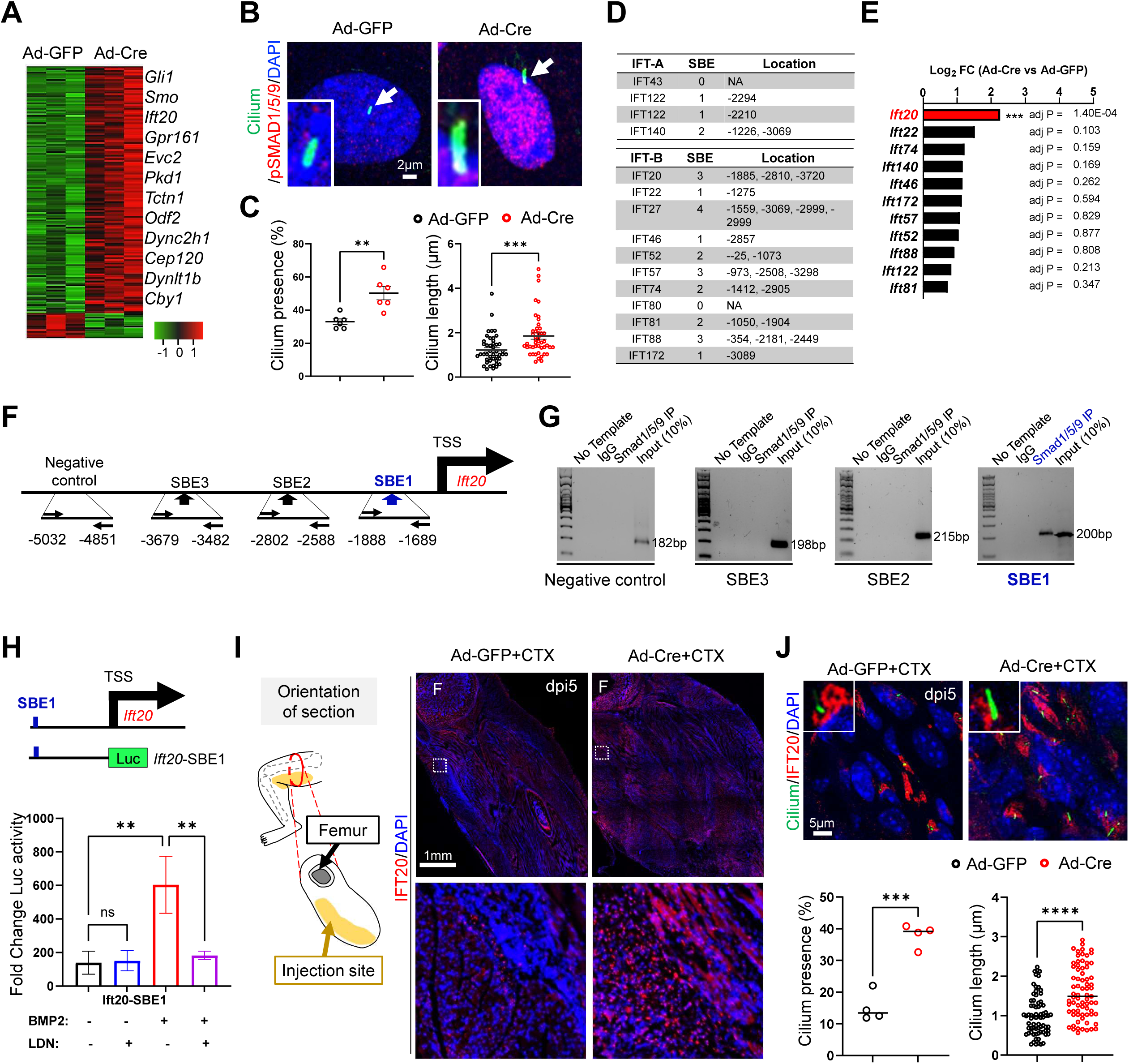
Enhanced BMP signaling induces elongation of primary cilia. **(A)** The heat map of RNA-seq shows the upregulated ciliary genes at days post-injury (dpi) 5 (n=3, p<0.05; t-test). **(B)** Subcutaneous mesenchymal progenitors (SMPs) isolated from *Acvr1^Q207D^* mice were stained with anti-acetylated tubulin and anti-phospho-SMAD1/5/9 antibodies. Arrows indicate cilia. **(C)** Cilia presence and length were quantified in Ad-GFR- or Ad-Cre-treated SMPs (n=3, **p<0.01, ***p<0.001; t-test). **(D)** The tables show the number and location of the Smad binding element (SBE) in IFT promoters. **(E)** The expression of *Ift* genes in SMPs was examined by qRT-PCR (n=3, ***p<0.001; t-test). **(F)** The location of SBEs (arrows) in the *Ift20* promoter. TSS: transcription start site. **(G)** PCR amplicons from ChIP analysis. **(H)** The upper panel shows the construct of the Ift20 luciferase (Luc) vector. The lower panel shows the result of the luciferase assay (n=3, **p<0.01; one-way ANOVA) treated with BMP2 ligand and BMP type I receptor inhibitor, LDN-193189 (LDN). **(I)** Ectopic IFT20 expression (red) is observed in the intramuscular damaged area (yellow highlighted in the left schema) at dpi5. Lower panels show higher magnification images from the boxes in the upper panels. F: femur. **(J) I**mmunohistochemistry images stained with anti-acetylated tubulin and IFT20 at dpi5. Cilia presence and length were quantified (n=4, ***p<0.001, ****p<0.0001; t-test).

Because primary cilium elongation depends on intraflagellar transport (IFT) complex dynamics (17, 34), we screened the promoter regions of the upregulated IFT transcripts for canonical Smad Binding Elements (SBE; 5′-CAGACA-3′), critical for transcribing BMP target genes (35, 36). *In silico* sequence analysis identified three conserved SBE sites within the proximal promoter region of *Ift20* (Figure 1D), a core intraflagellar transport component that emerged as a top candidate from our transcriptomic dataset (2.53-fold increase), confirmed via quantitative RT-PCR (Figure 1E). To establish physical interactions between the BMP transcriptional machinery and the *Ift20* locus, chromatin immunoprecipitation (ChIP) assays were performed on mesenchymal progenitors from the mouse embryonic limbs using an antibody specific for phosphorylated Smad1/5/9. Enriched transcription factor binding was detected explicitly at the SBE1 locus located within the proximal promoter region of *Ift20* (Figure 1F, G).

To confirm that this binding event directly activates transcription, we cloned the *Ift20* promoter region containing the SBE1 motif into a luciferase reporter plasmid. Recombinant BMP2 stimulation of transfected HEK293 cells significantly induced *Ift20* promoter-driven luciferase activity, an effect that was completely abolished by co-treatment with the selective BMP type I receptor inhibitor LDN-193189 (Figure 1H). *In vivo* immunohistochemical analysis confirmed that IFT20 expression was significantly elevated within the injury zone of Ad-Cre/CTX-injected *Acvr1^Q207D/+^*mice, correlating directly with the presence of elongated primary cilia (Figure 1I, J). Collectively, these data demonstrate that aberrant BMP signaling transactivates *Ift20* expression via canonical Smad complex binding to drive ciliary elongation during early HO pathogenesis.

### Ciliary elongation coordinates a dual-phase signaling relay essential for HO progression

Because the primary cilium is required for canonical Hh signal transduction (37–40), we investigated whether BMP-mediated ciliary lengthening renders progenitor cells hyper-responsive to extracellular Hh morphogens. Upon stimulation with the small-molecule Hh agonist SAG, the rate of ciliary accumulation of the Smo (40, 41) was markedly accelerated in Ad-Cre-treated *Acvr1^Q207D/+^* SMPs relative to control cells (Figure 2A, B). This rapid ciliary receptor sorting was accompanied by a synergistic upregulation of the definitive downstream Hh target gene *Gli1* (Figure 2C).

**Figure 2.**
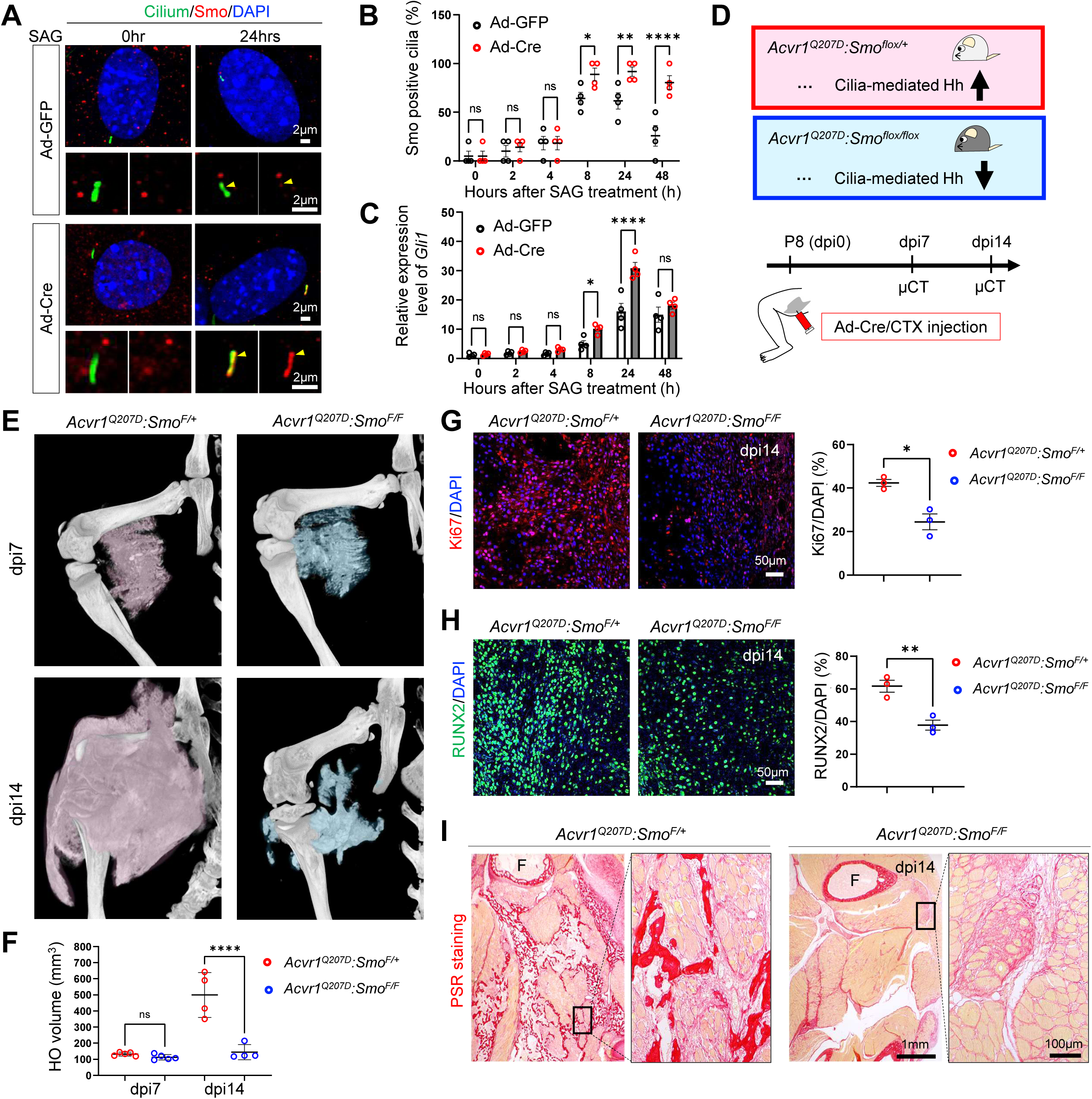
Suppression of cilia-mediated Hh signaling inhibits the late stage of HO development. **(A)** Smo (red) localization on cilia (green) was examined by immunocytochemistry 24 hours after smoothened agonist (SAG, 100nM) treatment, using Subcutaneous mesenchymal progenitors (SMPs) isolated from *Acvr1^Q207D^* mice. Yellow arrowheads indicate Smo accumulation on the cilia. **(B)** The percentage of Smo-positive cilia in SMPs was quantified at 0, 2, 4, 8, 24, and 48 hours after SAG treatment (n=4, *p>0.05, **p<0.01, ***p<0.001; t-test). **(C)** Gli1 expression level in SMPs was examined by qRT-PCR (n=4, *p<0.05, ****p<0.0001; t-test). **(D)** A genetic strategy to locally suppress cilia-mediated Hh signaling in *Acvr1^Q207D^*mice. Ad-Cre: adenovirus-Cre, CTX: cardiotoxin. **(E)** Ectopic bone formation was examined by micro-CT(µCT) at days post-injury (dpi) 7 and 14. **(F)** The volume of ectopic bone was measured (n=4, ****p<0.0001; t-test). **(G, H)** Proliferation and osteogenic differentiation are examined by staining with anti-Ki67 and anti-RUNX2 antibodies at dpi14. (n=3, *p<0.05, **p<0.01; t-test) **(I)** Picrosirius red (PSR) staining in dpi14. F: femur.

To establish the functional requirement of this ciliary signaling crosstalk *in vivo*, we crossed *Acvr1^Q207D/+^* mice with *Smo* floxed mice (*Smo^F/F^*) to achieve conditional, localized disruption of Hh transduction concurrent with HO induction (Figure 2D). Local administrative injury via Ad-Cre/CTX delivery did not alter ciliary lengthening or assembly at dpi7 in *Acvr1^Q207D/+^:Smo^F/F^* mice (Figure S2A, B). Longitudinal µCT imaging revealed a striking temporal divergence: conditional ablation of *Smo* did not block the initial tissue condemnation or the onset of HO between dpi0 and dpi7. However, the subsequent volumetric expansion of mature heterotopic bone mass was profoundly attenuated in *Acvr1^Q207D/+^:Smo^F/F^* mice between dpi7 and dpi14 (Figure 2E, F). Histological evaluations confirmed that local *Smo* deletion suppressed downstream ectopic proliferation, osteogenic maturation, and mature osteoblast matrix deposition (Figure 2G-I). These findings demonstrate that while the ciliary-Hh pathway is dispensable for the initial fate transition and tissue condemnation phase of HO, but required during a late phase to drive the proliferative expansion and maturation of the HO.

### Complete structural ablation of the cilium phenocopies and exceeds Hh blockade

The temporal divergence observed in our *Smo*-deficient mice led us to hypothesize that the primary cilium coordinates a multi-stage signaling relay, where the physical presence of the ciliary axoneme is required for early, Hh-independent lineage commitment, while ciliary Hh signaling sequentially drives later tissue expansion (Figure 2). To test whether complete structural ablation of the cilia would yield a greater rescue than *Smo* deletion alone, we deployed both pharmacological and specific genetic strategies to disrupt ciliary assembly. *In vitro*, treatment of isolated *Acvr1^Q207D/+^*SMPs with the ciliary assembly blocker Ciliobrevin A (CiA) (42, 43) effectively neutralized the accelerated ciliogenesis induced by Ad-Cre expression (Figure 3A, B). Without CiA treatment, osteogenic differentiation was enhanced in Ad-Cre-treated SMPs compared with Ad-GFP-treated SMPs (Figure 3C). For *in vivo* translation, *Acvr1^Q207D/+^* mouse pups were injected with Ad-Cre/CTX at postnatal day 8 (P8) and treated systemically with CiA (5 mg/kg per day) from P9 to P14 (Figure 3D). Longitudinal µCT analysis demonstrated that pharmacological disruption of early ciliary assembly induced a dramatic reduction in mature heterotopic bone volume (Figure 3E, F) without altering systemic skeletal density or total body weight, effectively phenocopying the genetic ablation of ciliogenesis (Figure S3A, B).

**Figure 3.**
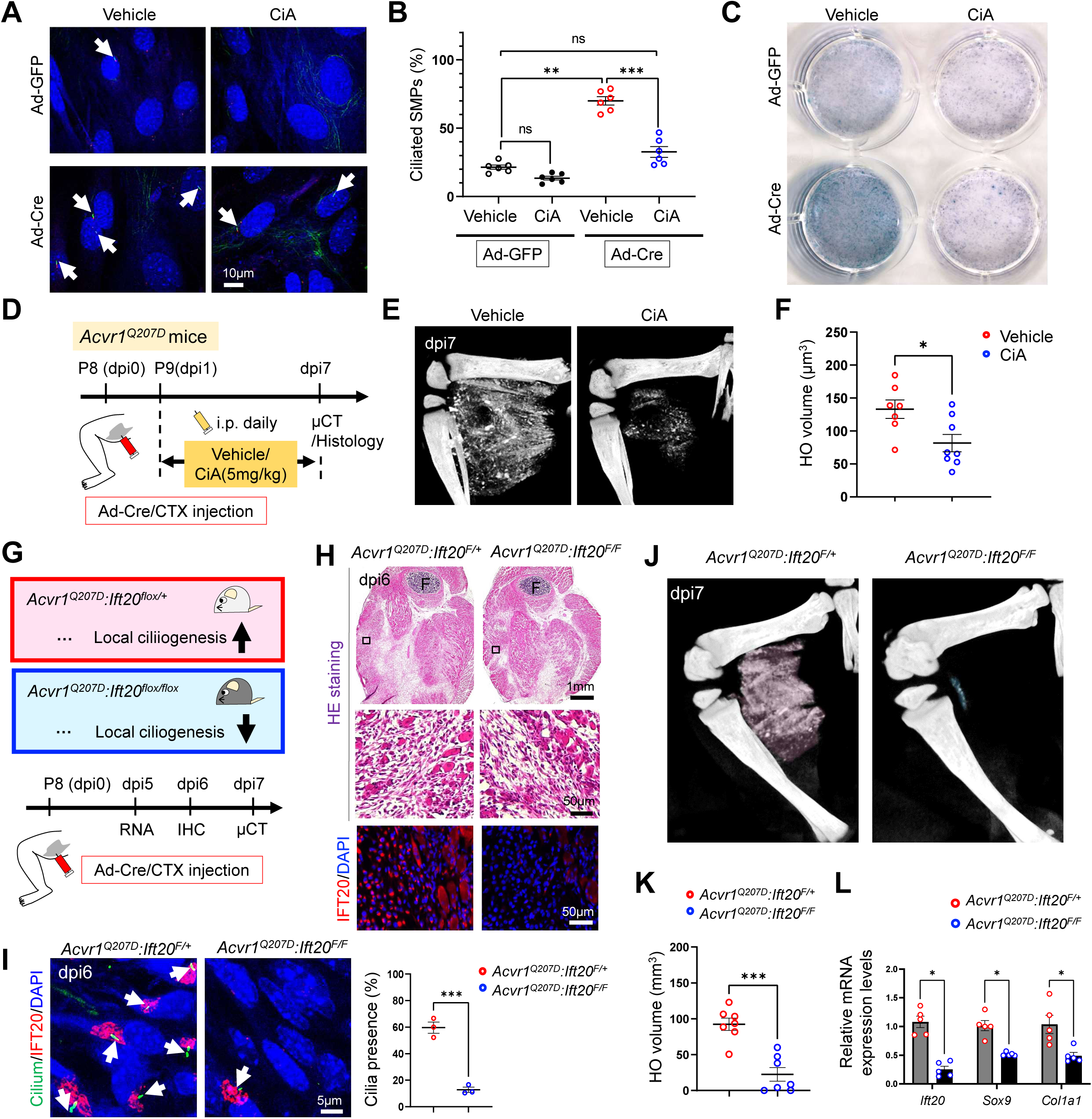
Pharmacological and genetic suppressions of ciliogenesis inhibit HO development. **(A)** Ad-GFP and Ad-Cre transduced subcutaneous mesenchymal progenitors (SMPs), isolated from *Acvr1^Q207D^* mice, were treated with either vehicle or 50µM ciliobrevin-A (CiA). Arrows show cilia stained with anti-acetylated-tubulin and gamma-tubulin antibodies. **(B)** Quantification of ciliated SMPs (n=6, **p<0.01, ***p<0.001; one-way ANOVA). **(C)** Alkaline phosphatase staining of SMPs 3 days after culturing with osteogenic medium. **(D)** Schematic schedule for CiA treatment to *Acvr1^Q207D^* mice. **(E)** Representative µCT images at days post-injury (dpi)7. **(F)** Quantification of the HO volume (n=8, *p<0.05; t-test). **(G)** A genetic strategy to locally suppress cilia in *Acvr1^Q207D^* mice. **(H, I)** Cilia (green) and IFT20 (magenta) were examined by immunohistochemistry at dpi6. The presence of cilia in damaged muscle area was quantified (n=3, ***p<0.001; t-test). F: femur. **(J)** Representative µCT images at dpi7. **(K)** Quantification of the HO volume (n=7, ***p<0.001; t-test). **(L)** mRNA expression level was examined by qRT-PCR (n=5, *p<0.05; t-test).

To definitively isolate this structural requirement without chemical confounding, we crossed *Acvr1^Q207D/+^* mice with *Ift20* floxed mice (*Ift20^F/F^*) to generate *Acvr1^Q207D/+^:Ift20^F/F^* mice to achieve conditional deletion of the essential ciliary transport component *Ift20* within the injury microenvironment. Local delivery of Ad-Cre simultaneously activated pathogenic ACVR1 signaling while completely disrupting structural ciliary assembly (Figure 3G). We found that ectopic IFT20 expression coincided with the presence of cilia in HO mice (Figure 3H), while the local deletion of *Ift20* suppressed ciliogenesis and inhibited HO in rescue mice (Figure 3I-K). Critically, unlike the *Smo* knockout models, which only halted late-stage progression, genetic ablation of the ciliary structure completely suppressed HO development at its inception, reducing the transcription of foundational osteochondrogenic commitment markers, including *Sox9* and *Col1a1*, as early as dpi5 (Figure 3L). These data demonstrate a hierarchical epistatic requirement for the primary cilium: the physical organelle is required for early pro-osteochondrogenic lineage commitment through an Hh-independent mechanism, whereas the ciliary Hh signaling is subsequently utilized to drive HO expansion.

### The BMP-IFT20-Cilia axis is conserved in FOP mouse models driven by the authentic *Acvr1^R206H^* mutation

To establish the translational relevance of this multi-stage ciliary mechanism to human disease, we examined whether ciliary remodeling operates in the context of the classical Acvr1^R206H^ mutation that accounts for the vast majority of clinical FOP cases. Re-analysis of a public transcriptomic dataset from an *Acvr1^R206H/+^* FOP mouse model (44) revealed a profound, localized enrichment of ciliary assembly and intraflagellar transport pathways within active HO lesions, highlighting a significant upregulation of *Ift20* (Figure 4A, B).

**Figure 4.**
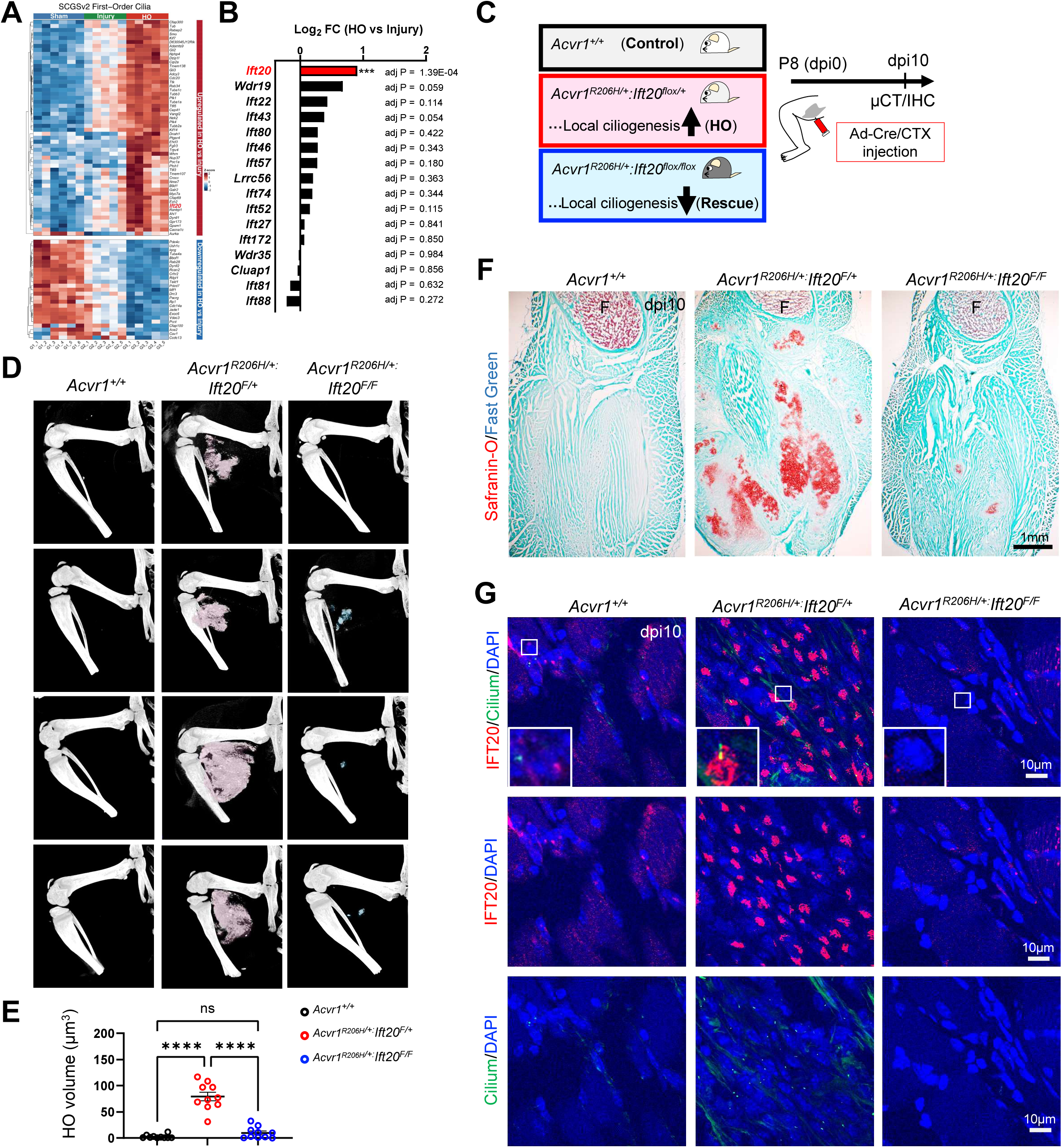
The BMP-IFT20-Cilia axis is conserved in FOP mouse models. **(A, B)** The heatmap using a public transcriptomic dataset from an *Acvr1^R206H/+^* FOP mouse model, highlighting a significant upregulation of *Ift20* in the FOP mouse model. **(C)** A genetic strategy to locally suppress cilia in *Acvr1^R206H^*FOP mice. **(D)** Representative µCT images at days post-injury (dpi) 10. **(E)** Quantification of the HO volume (n=10, ****p<0.0001; one-way ANOVA). **(F)** Safranin-O staining at dpi10. F: femur. **(G)** Cilia (green) and IFT20 (magenta) were examined by immunohistochemistry at dpi10.

To test whether targeted ciliary disruption could rescue the authentic FOP phenotype, we crossed *Acvr1^[R206H]FlEx/+^* mice (6) and *Ift20* flox mice to generate conditional *Acvr1^[R206H]FlEx/+^:Ift20^F/F^* mice (Figure 4C). Following local administrative injury via Ad-Cre/CTX, control *Acvr1^[R206H]FlEx/+^:Ift20^F/+^*mice developed extensive HO by dpi14 (Figure 4D; middle panels, 4E). Conversely, homozygous conditional deletion of *Ift20* in *Acvr1^[R206H]FlEx/+^:Ift20^F/F^* mice robustly mitigated ectopic ossification volume (Figure 4D, right panels). Histological profiling demonstrated that ciliary ablation via *Ift20* knockout dramatically impaired the preliminary mesenchymal condensation and chondrogenesis phases (Figure 4F), matching a complete loss of localized IFT20 expression and ciliary structures within the soft-tissue lesion (Figure S4, 4G).

### Pathological BMP signaling reprograms fibro-adipogenic progenitor ciliogenesis to develop HO

Finally, we wished to clarify the specific cell populations undergoing this ciliary-mediated remodeling within the injury microenvironment. Given that PDGFRα-positive FAPs represent a major cellular origin of heterotopic bone following tissue trauma (7, 45–48), we examined temporal ciliary dynamics specifically within this population *in vivo*. In control tissue (Ad-GFP/CTX-treated *Acvr1^Q207D/+^* mice), ciliated FAPs expanded transiently to a peak at dpi5 before returning to baseline levels by dpi14 in *Acvr1^Q207D/+^* mice (Figure S5, 5A). In contrast, FAPs expressing pathogenic Ad-Cre/CTX-treated *Acvr1^Q207D/+^* mice exhibited sustained, non-resolving increases in total ciliation frequency alongside profound structural axoneme lengthening *in vivo* (Figure 5B). To confirm that this ciliary remodeling within FAPs drives the authentic FOP phenotype, we assessed FAP behavior within Ad-Cre/CTX-treated *Acvr1^[R206H]FlEx/+^* mice. Pathogenic *Acvr1^[R206H]FlEx/+^* expression led to a massive expansion of hyper-ciliated FAP populations within the intermuscular regions (Figure S6, 5C, middle panels). However, conditional co-deletion of *Ift20* efficiently suppressed this aberrant ciliary expansion (Figure 5C, right panels). Quantification analysis confirmed that ciliogenesis was robustly attenuated in conditional co-deletion of *Ift20* in *Acvr1^[R206H]FlEx/+^* mice (Figure 5D). At the molecular level, FAPs lacking *Ift20* failed to upregulate the Hh target transcript *Gli1* (Figure 5E, S7), leading to a significant reduction in local mesenchymal proliferation rates within the early lesion (Figure 5F). These data demonstrate that both hyperactivated and pathological mutation-driven ACVR1 signaling structurally reprogram the primary cilium that governs the pathogenesis of HO (Figure 5G).

**Figure 5.**
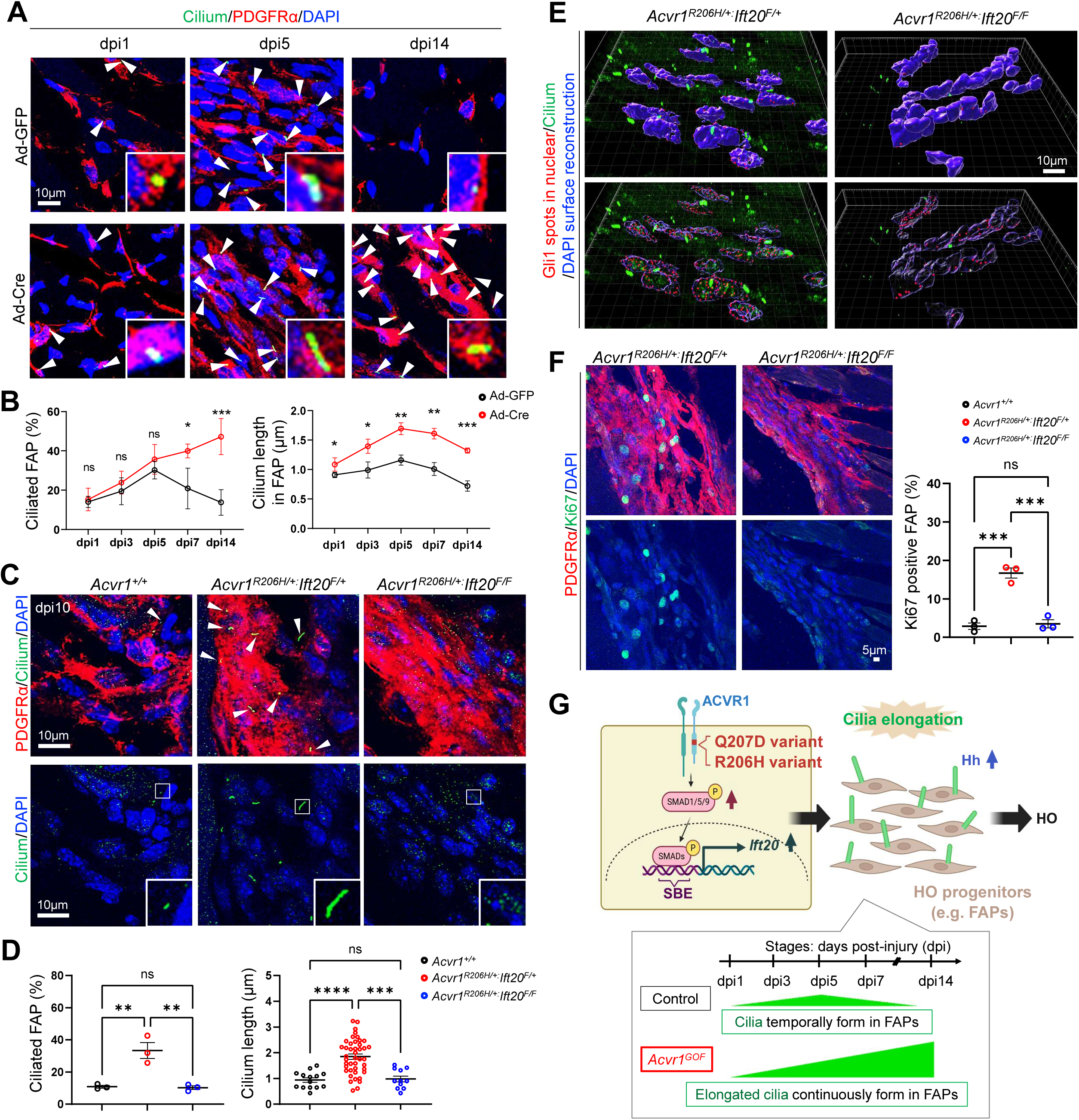
Pathological BMP signaling reprograms fibro-adipogenic progenitor ciliogenesis to develop HO. **(A)** Ciliated fibro-adipogenic progenitors (FAPs) (arrowheads) in Ad-GFP and Ad-Cre treated *Acvr1^Q207D^* mice were examined by immunohistochemistry with anti-PDGFRα and anti-ARL13B antibodies at days post-injury (dpi) 1, 5, and 14. **(B)** The presence and length of cilia in FAPs were quantified (n=3, *p<0.05, **p<0.01, ***p<0.001; t-test). **(C)** Ciliated FAPs (arrowheads) in *Acvr1^R206H^* mice were examined by immunohistochemistry with anti-PDGFRα, and anti-ARL13B antibodies at dpi 10. **(D)** The presence and length of cilia in FAPs were quantified (n=3, **p<0.01, ***p<0.001, ****p<0.0001; one-way ANOVA). **(E)** Nuclear (Blue reconstructed-surface) localization of Gli1 (Red spots), and presence of cilia (green) were examined by Imaris 10.2.0 software. **(F)** The number of Ki67-positive FAPs was examined by immunohistochemistry (n=3, ***p<0.001; one-way ANOVA). **(G)** Both hyperactivated (Q207D)- and pathological mutation (R206H)-driven ACVR1 signaling induces the elongation and continuous presence of primary cilia in HO progenitors (e.g. FAPs) through transcriptional regulation of *Ift20*, which governs the pathogenesis of HO.

## Discussion

Aberrant activation of BMP signaling through mutations in the type I receptor ACVR1 is the defining driver of FOP and a critical catalyst in HO. While the downstream transcriptional responses of pathogenic ACVR1 signaling have been extensively studied, the structural and organelle-level remodeling required to sustain these osteogenic signals in the injury microenvironment remains poorly defined. Here, we demonstrate that hyperactivated or pathological BMP signaling drives ectopic bone formation by structurally and functionally reprogramming the primary cilium in HO progenitors such as FAPs. By linking canonical Smad1/5/9 transcriptional machinery directly to the transactivation of the IFT component, our findings reveal an active growth factor-organelle signaling loop that governs HO pathogenesis.

### Pathogenic BMP signaling is a critical driver to enhance ciliogenesis in HO development

Our previous work established that genetic loss of *Acvr1* impairs mouse embryonic node ciliogenesis via cell cycle attenuation (29, 30); conversely, the data presented here demonstrate that pathological activation of ACVR1 expands the ciliary axoneme and increases ciliation frequency in a cell-autonomous manner in HO development. Using stem cells from human exfoliated deciduous teeth (SHEDs) from FOP patients, a recent study showed subtle differences in ciliogenesis, i.e., shorter cilia length in FOP-SHEDs compared with wild-type SHEDs (49). This discrepancy can be explained by the examined cell types in different HO progenitors. For example, expression levels of numerous ciliary genes were significantly increased in FOP tissues, including several HO progenitors in mice (Figure S3A). Expression levels of IFT genes are also highly increased in tendon progenitors in trauma-induced HO in mice (50). In agreement with these findings, we found that structural cilia elongation is driven by direct binding of phosphorylated Smad1/5/9 to an SBE1 within the proximal promoter of *Ift20* (Figure 1F-H). While there were several SBE sites found in the putative promoter region in other IFTs, enhanced transcription of *Ift20* seems to be responsible for the induction of HO. This notion is further supported by the analysis of a public transcriptomic dataset from an FOP mouse model, showing that among IFT components, *Ift20* expression is statistically increased (Figure 4A, B). Therefore, our study establishes a new link between BMP pathways and cilia biogenesis that is critical to understanding the novel pathogenesis of HO, mainly driven by IFT20-associated ciliogenesis.

### Primary cilium may operate via a dual-phase signaling relay to develop HO

By parallel evaluation of structural ciliary ablation (i.e., genetic *Ift20* deletion or pharmacological Ciliobrevin A treatment) (Figure 3E, K) and ciliary signal transduction blockade (i.e., genetic *Smo* deletion) (Figure 2E), we uncovered a strict hierarchical epistatic requirement for the primary cilium that operates via a dual-phase signaling relay. During the early HO lineage commitment phase, complete structural ablation of the cilium completely suppresses the induction of foundational osteochondrogenic transcripts such as *Sox9* and *Col1a1* (Figure 3L). Strikingly, conditional deletion of *Smo* has no impact on this early tissue condemnation phase. This demonstrates that the physical presence of the ciliary structure regulates initial pro-osteochondrogenic lineage commitment through a non-canonical, Hh-independent mechanism. This early window may rely on the compartmentalization or trafficking of alternative ciliary-localized pathways, such as TGF-β or PDGF clustered at the ciliary base (51, 52). During the HO expansion phase, the elongated ciliary axoneme may act as an amplifier for canonical Hh signaling. Elongated cilia exhibit accelerated ciliary sorting and accumulation of Smo, driving synergistic transactivation of *Gli1*. While dispensable for initial fate transitions, this ciliary Hh relay may be required to drive the subsequent proliferation of HO progenitors and osteogenic maturation necessary for the volumetric expansion of mature HO (Figure 2E). While there is numerous evidence showing that Hh signaling is the key regulator to develop HO (32, 33, 53), this study may provide the resolution of a long-standing ambiguity regarding the timing and necessity of Hh signaling during HO development.

### Altered cilia remodeling in FAPs can be a novel cellular mechanism of HO in FOP mice

Our *in vivo* tracking identifies PDGFRα-positive FAPs as the principal cell population undergoing this pathological ciliary remodeling. Physiologically, resolving wound healing, ciliated FAP populations expand transiently and rapidly return to baseline (54). In wild-type mice, under the CTX-induced muscle injury condition, the proportion of PDGFRα-positive FAPs with cilia increases sharply from 10% to almost 50% before returning to pre-injury levels at 21dpi (54). Under the influence of hyperactivated BMP signaling via *Acvr1^Q207D/+^* or pathogenic *Acvr1^R206H/+^* mutations, however, injury-primed FAPs display a sustained, non-resolving hyper-ciliation state (Figure 5A-E). This persistent morphological defect may lock FAPs into a proliferative, osteogenic loop, transforming a transient regenerative cell type into osteochondroprogenitors. Importantly, the conservation of the BMP-IFT20-Cilia axis in the authentic *Acvr1^R206H/+^* mouse model establishes the direct translational relevance of these findings to human FOP. While our study does not focus on muscle regeneration, during muscle repair, FAPs do not directly undergo differentiation into myofibers (46–48); FAPs support the self-renewal, proliferation, and differentiation of muscle stem cells (MuSCs) (55, 56). Since inhibiting ciliogenesis in FAPs enhances myofiber regeneration in mice (54) and FAPs, but not MuSCs, are involved in defective myogenic activity in FOP mice (57), elongated cilia in FAPs may inhibit MuSCs differentiation by modulating paracrine mechanisms, thereby attenuating muscle regeneration in FOP mice. We will address this attractive question in the future.

Although our study provides proof of principle that BMP signaling drives HO by structurally stimulating the primary cilia, it has several limitations. First, while the cilia-mediated Hh dual-phase model demonstrates that early pro-osteochondrogenic lineage commitment requires the physical ciliary structure but is independent of Hh signaling, the alternative ciliary pathways remain speculative. Second, since we used Ad-Cre/CTX to induce HO, this hyper-inflammatory model may not faithfully recapitulate human FOP, which is frequently characterized by spontaneous, non-injury-driven flare-ups. Third, systemic delivery of the ciliary blocker Ciliobrevin A effectively mitigated HO. However, because primary cilia are ubiquitous organelles, the lack of long-term toxicity profiles limits their immediate clinical translatability. Lastly, we wished to focus almost exclusively on cell-autonomous ciliary remodeling within PDGFRα-positive FAPs. However, it remains unknown how activated BMP signaling structurally modifies the cilia in FAPs without FAP-specific deletion of cilia in the HO model. This experiment will be critical to identify other critical niche cells (e.g., endothelial or immune cells) or to understand how ciliary ablation alters FAP-mediated paracrine signaling in HO.

In conclusion, our study demonstrates that BMP signaling acts as a direct upstream transcriptional regulator of ciliogenesis, establishing a novel mechanism critical for HO development. Because the structural ciliary abnormalities are restricted to the active injury zone, targeting ciliary assembly or its specific downstream IFT dynamics may offer a temporally targetable strategy to arrest HO, bypassing the limitations of systemic treatment with pharmacological inhibitors.

## Experimental Procedures

### Animals

*Acvr1^Q207D/+^* mice (28) and *Acvr1^[R206H]FlEx/+^* mice (6) were obtained from Dr. Yuji Mishina and Dr. Aris Economides, respectively. *Smo* floxed mice (58) and *Ift20* floxed mice (59) were obtained from the Jackson Laboratory. To develop HO in *Acvr1^Q207D/+^* mice and *Acvr1^R206H/+^* mice, Ad-GFP or Ad-Cre from Vector Development Lab (∼0.5 × 10^9^ PFU per mouse, Baylor College of Medicine) and cobra venom factor (0.03 μg per mouse, Sigma, 217503) were co-injected into the right limbs at postnatal day 8. After Ad-Cre/CTX injection into *Acvr1^Q207D/+^* mice, mice were injected intraperitoneally with Ciliobrevin A (Sigma, H4541) at 5 mg/kg (n=8) or vehicle (n=7) daily for 6 days. All mice were maintained in the animal facility of the McGovern Medical School at UTHealth Houston. The experimental protocol was reviewed and approved by the Animal Welfare Committee and the Institutional Animal Care and Use Committee of UTHealth Houston.

### Histological analysis, immunofluorescent staining, and micro-computed tomography (μCT) analysis

H&E staining, Alkaline phosphatase staining, Safranin O staining, and immunofluorescent staining were performed as described previously (60, 61). Picrosirius red staining was performed using 1% picrosirius red solution (Sigma-Aldrich, 365548 and P6744). Primary antibodies used in immunofluorescent staining were as follows: acetylated tubulin (1:1000, Sigma-Aldrich, T6793), ARL13B (1:200, Proteintech, 17711-1-AP), gamma-tubulin (1:1000, Sigma-Aldrich, T5326), GLI1 (1:200, R&D system, AF3455), IFT20 (1:250, Proteintech, 13615-1-AP), Ki-67 (1:500, BD, 550609), PDGFRα (1:100, R&D, AF1062), and RUNX2 (1:250, Cell Signaling, #12556). Slides were viewed with an Olympus FluoView FV1000 laser scanning confocal microscope using the FV10-ASW Viewer (Ver. 3.1). Quantification of immunofluorescent signals was analyzed by IMARIS software (version 10.2.0). Micro-computed tomography (μCT) images were scanned with a CT system at 50 kV energy and 200 μA intensity (Skyscan 1276; Bruker, Billerica, MA). The slices under ectopic bone were reconstructed to produce 3D images, and bone volume was measured using CTAn (Bruker, Billerica, MA.).

### Isolation and culture of subcutaneous mesenchymal progenitor cells (SMPs)

SMP cells were isolated and cultured by following the methods previously (33). Briefly, mouse adipose deposits were removed under sterile conditions, and tissue was minced and digested with collagenase type I (1 mg/ml) and 0.5% trypsin in 0.1% bovine serum albumin for 70 min at 37°C. Digested tissue was centrifuged at 500g for 10 min, and the pellet was carefully collected after aspirating off the floating fat depots. After a second centrifugation at 500g for 10 min, the cellular pellet was filtered through a 100-μm mesh filter to remove debris.

### Immunocytochemistry

SMP cells were fixed with 4% paraformaldehyde for 10 minutes at room temperature (RT) and permeabilized with 0.1% Triton X-100 in PBS (PBST) for 15 minutes at RT. They were then blocked in 2% sheep serum in PBST for 30 minutes at RT prior to staining with primary antibodies. The primary antibodies used were as follows: acetylated tubulin (1:2000, Sigma-Aldrich, T6793), gamma-tubulin (1:2000, Sigma-Aldrich, T5326), IFT20 (1:500, Proteintech, 13615-1-AP), and phospho-Smad1/5/9 (1:250, Cell Signaling, 13820), and smoothened (1:400, Abcam, ab38686). Images were captured with an Olympus FluoView FV1000 laser scanning confocal microscope equipped with the software FV10-ASW Viewer (Ver. 3.1), and Zeiss Axio Zoom equipped with the ZEN microscopy software.

### Quantitative RT-PCR

Using Trizol Reagent (Thermo Fisher Scientific), total RNA was extracted from hindlimb tissues or from SMP cells. The total RNA was treated with DNase I (Roche) before cDNA synthesis. cDNA was synthesized using iScript Reverse Transcription Supermix for RT-qPCR (Bio-Rad). Quantitative RT-PCR was carried out using SsoAdvanced Universal SYBR Green Supermix (Bio-Rad). The sequences of the specific primer sets are listed in Supplemental Table 1.

### Luciferase assays

HEK293 cells were cultured in DMEM supplemented with 10% FBS (Fisher) and penicillin/streptomycin and transfected by FuGENE HD^®^ (Promega). *pGL3-Ift20-SBE-1* transfected cells were incubated in 60 mm culture dishes. After 24 hours of incubation, the medium was removed, then serum-free DMEM was added for serum starvation for 24 hours. After serum starvation, recombinant human BMP2 protein (50 ng/ml, R&D, 355-BM) and LDN193189 (0.5 µM, Millipore Sigma, SML0559) were treated. After 6 hours of incubation, reporter activities were measured by a luminometer. All luciferase activities were normalized to β-galactosidase activity and are shown as mean-fold differences relative to empty luciferase plasmids. The pGL3 basic empty vector was added to each experiment as a mock control. CMV β-galactosidase reporter plasmids were co-transfected in all experiments as a control for transfection efficiency.

### Chromatin immunoprecipitation (ChIP) Assay

The ChIP assays were performed by using the ChIP-IT^®^ Express Kit (ACTIVE MOTIF) with some modifications. Wild-type mouse hindlimbs at E11.5 were harvested, sonicated by using the S220 Covaris sonication instrument with Sonolab™ Software (Version7.1.4) and truChIP Chromatin Shearing Reagent Kit (Covaris Inc). Shearing efficiency was performed to check chromatin size before doing ChIP. Pierce Protein A Magnetic Beads (Thermo Fisher Scientific Inc) were blocked with BSA (10mg/ml) and herring sperm gDNA at 4°C overnight. ChIP reactions were set as follows: preclear chromatin with IgG, preclear chromatin with Smad1/5/9 (Cell Signaling, 13820) antibody, no chromatin (Smad1/5/9 antibody only). Chromatins were eluted with Reverse-crosslinking buffer and purified with phenol and chloroform. All PCR reactions were done under an annealing temperature of 60 °C. The sense primer (5’-CCAGTCGTAGGAGTTCTGAT-3’) and the antisense primer (5’-CTTGTCTTCTCTGGCTTCC -3’) were used to amplify the *Ift20* promoter containing the Smad1/5/9 binding element in *Ift20-SBE1*. All the PCR products were evaluated on a 2% agarose gel and confirmed by sequencing. As controls, the primers were used in PCR without chromatin; without antibody; normal rabbit IgG was used, replacing the Smad1/5/9 antibody to reveal nonspecific immunoprecipitation of the chromatin. Quantification of the Smad1/5/9-binding DNA fragment was performed using the same primers and conditions as above. Each reaction was done in triplicate, and S.E. was calculated from 3 independent experiments. Primers to a -5032 bp upstream of *Ift20* transcriptional start site of the Smad1/5/9 binding element were used as controls. (Sense: 5’-TCTAGGATCAGCTCTGCTTC-3’, antisense: 5’-CCTGAGACAGGAAGTGTATC-3’).

### Cilia gene reference

Ciliary genes were defined using the SYSCILIA Gold Standard version 2 (62), comprising 686 genes in three confidence tiers: first-order (high-confidence ciliary, 539 genes), second-order (132 genes), and others. Human gene symbols were mapped to mouse symbols using the homologene package, with title-case symbol matching as a fallback for entries without a homologene record. Genes mapping to symbols not detected in the bulk count matrix were excluded from downstream analyses.

### Bulk RNA-seq differential analysis

The bulk RNA-seq data were obtained from GEO (GSE220725), which comprises gastrocnemius muscle from FOP-ACVR1 transgenic mice subjected to a pinch-injury model and sampled at day 7 post-injury (44). Sixteen samples were retained for analysis across three groups: G1 (Sham; DOX–, pinch–; uninjured baseline; n=6), G2 (Injury; DOX–, pinch+; injury without FOP allele; n=5), and G3 (HO; DOX+, pinch+; injury with active FOP allele, HO-forming; n=5). FASTQ files were quantified against the GENCODE vM38 mouse transcriptome using Salmon v1.10 (63). Differential expression was assessed using DESeq2 v1.42 (64). P-values were adjusted using the Benjamini-Hochberg with significance defined as adjusted p < 0.05 with |log2 fold change| > 0.58.

### Statistical analysis

The Student’s *t*-test and one-way ANOVA were used for statistical analysis. A P-value of less than 0.05 was considered statistically significant.

## Author contributions

Y.K. designed the study. H.Y., J.W., F.Y., J.B., and Y.K. performed the experiments. H.Y., J.W., F.Y., J.B., Z.Z., and Y.K. analyzed the data. H.Y., R.D., W.L., A.E., Y.M., and Y.K. wrote the manuscript.

## Acknowledgements

We gratefully acknowledge Dr. Yingzi Yang for guidance on SMP cell isolation and culture, and Dr. Sean P. Marrelli for assistance with μCT analysis. We also thank Dr. Maurizio Pacifici, Dr. Dan Perrien, Dr. Rebecca Berdeaux, and Dr. Benjamin Levi for the insightful discussion. We also thank Dr. Adrian Romero Mora for technical assistance and Ms. Maryam Faisal for sample preparation. Micro-CT imaging was performed through the μCT Imaging Core Facility at the McGovern Medical School at UTHealth, which is supported by NIH S10OD030336 (S.M.). This work was partially supported by the McGovern Medical School at UTHealth Houston Retention Fund (Y.K.), a research grant NIDCR/NIH R01DE025897 (Y.K.), and by the Bone Disease Program of Texas Rolanette and Berdon Lawrence Research Award (H.Y.). Schematic illustrations were created by BioRender software (a license purchased by the University of Texas Health Science Center at Houston). We thank the support from the Cancer Prevention and Research Institute of Texas (CPRIT RP180734 and RP240610).

---

**Supplemental Figure 1.**
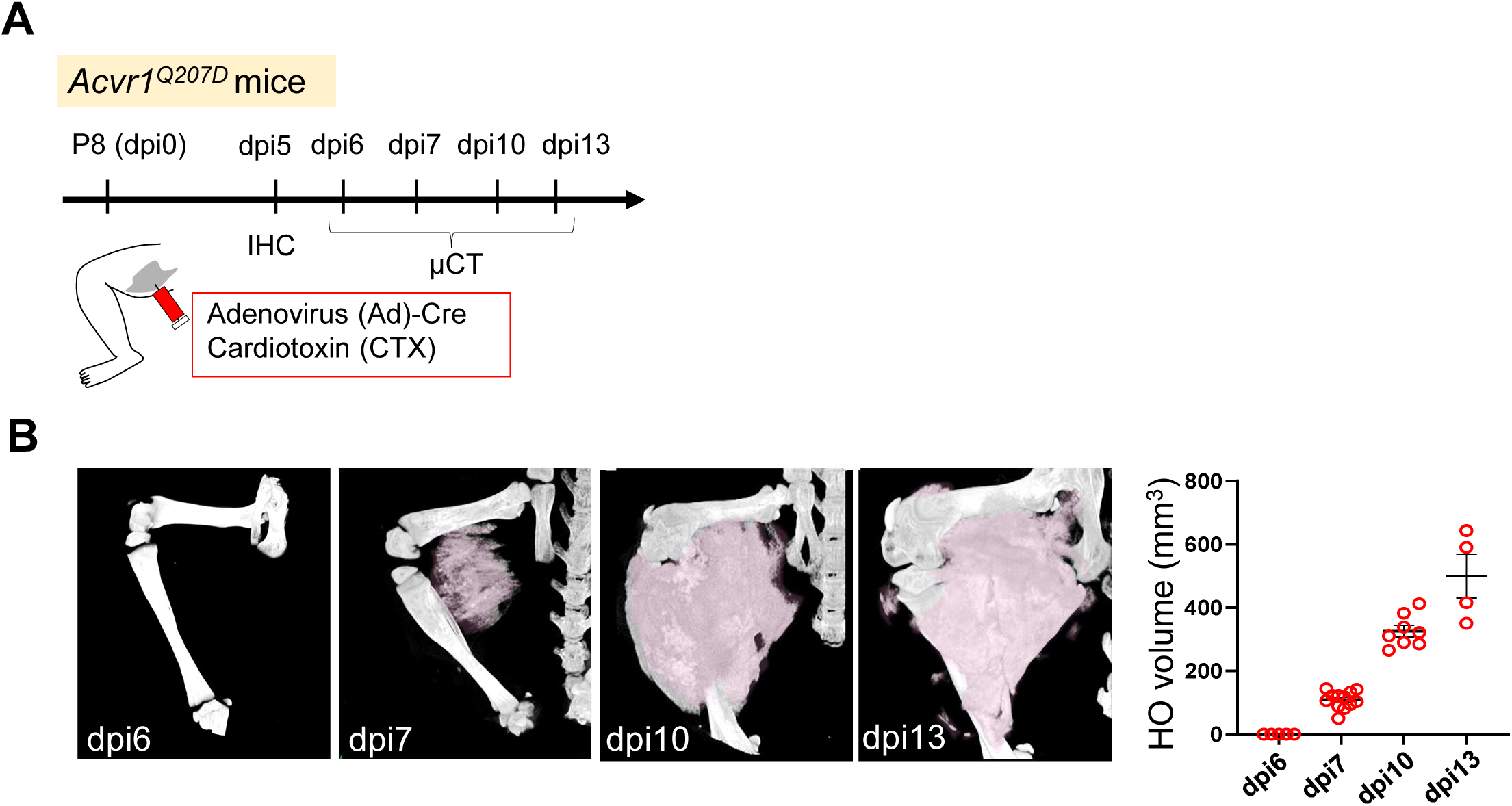
HO volume in *Acvr1^Q207D^* mice. **(A)** Experimental schedule. **(B)** Representative micro-CT images and quantification of HO volume at dpi6, 7, 10, and 13.

**Supplemental Figure 2.**
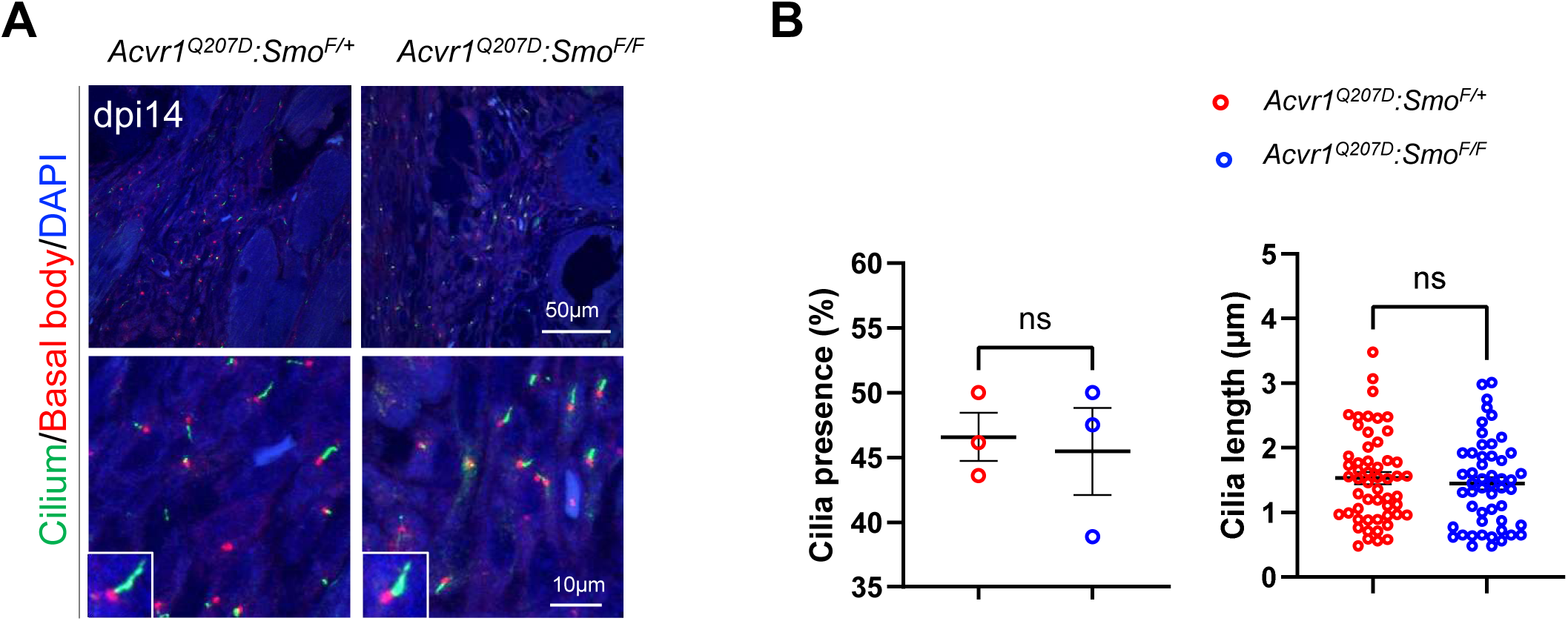
Local suppression of ciliary-mediated Hh signaling doesn’t affect cilia formation. **(A)** Immunohistochemistry with anti-acetylated tubulin and anti-gamma-tubulin antibodies at dpi14. Lower panels are high-magnification images from upper panels. **(B)** Quantification of cilia presence and length in the damaged area (n=3, ns: not significant, t-test).

**Supplemental Figure 3.**
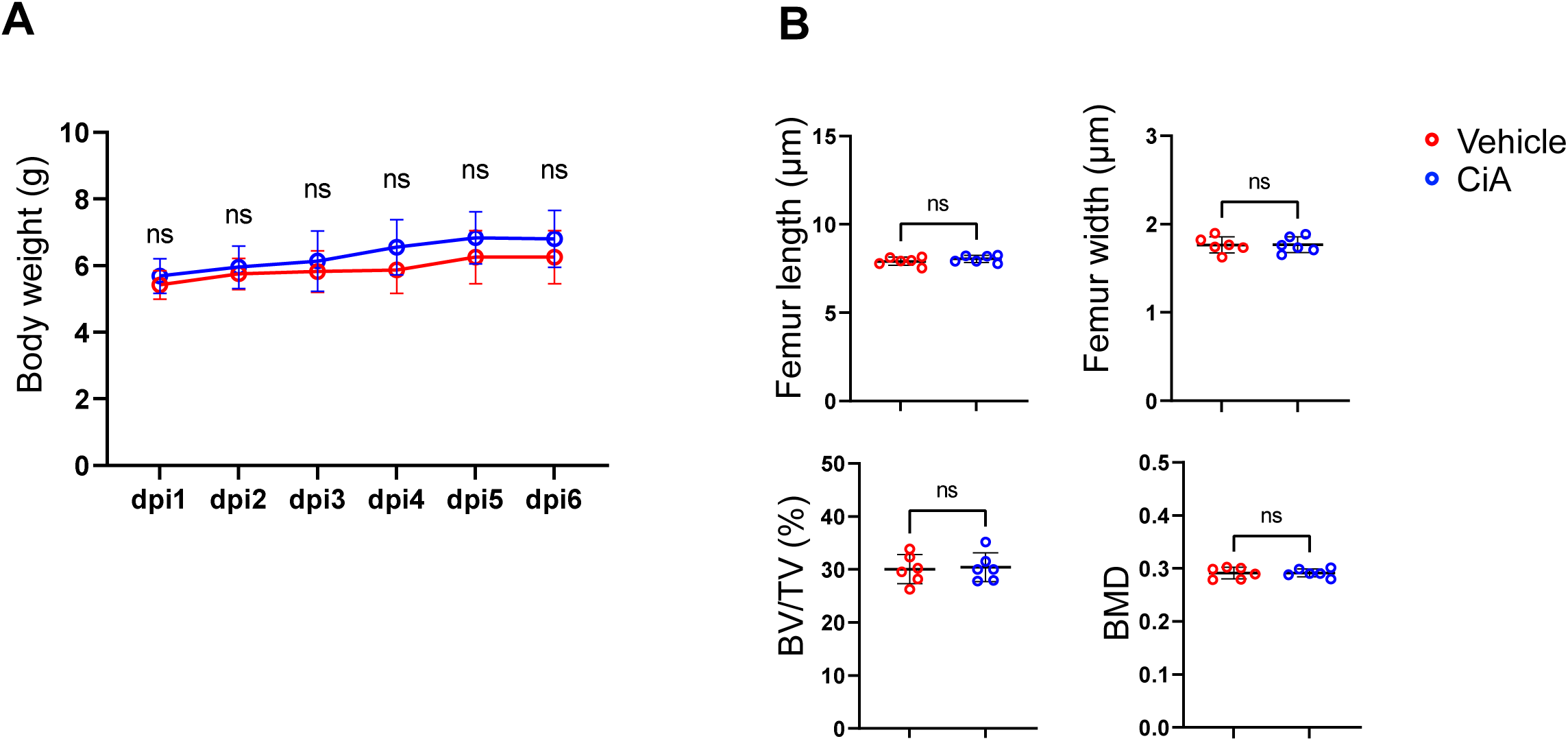
Ciliobrevin A treatment doesn’t cause bone loss or body weight loss. **(A)** Body weight during ciliobrevin A (CiA) treatment in *Acvr1^Q207D^* mice (n=6, ns: not significant, t-test). **(B)** Femur length, femur width, Bone volume (BV)/ Tissue volume (TV) in femur, and bone mineral density (BMD) after either vehicle or CiA treatment at dpi7 (n=6, ns: not significant, t-test).

**Supplemental Figure 4.**
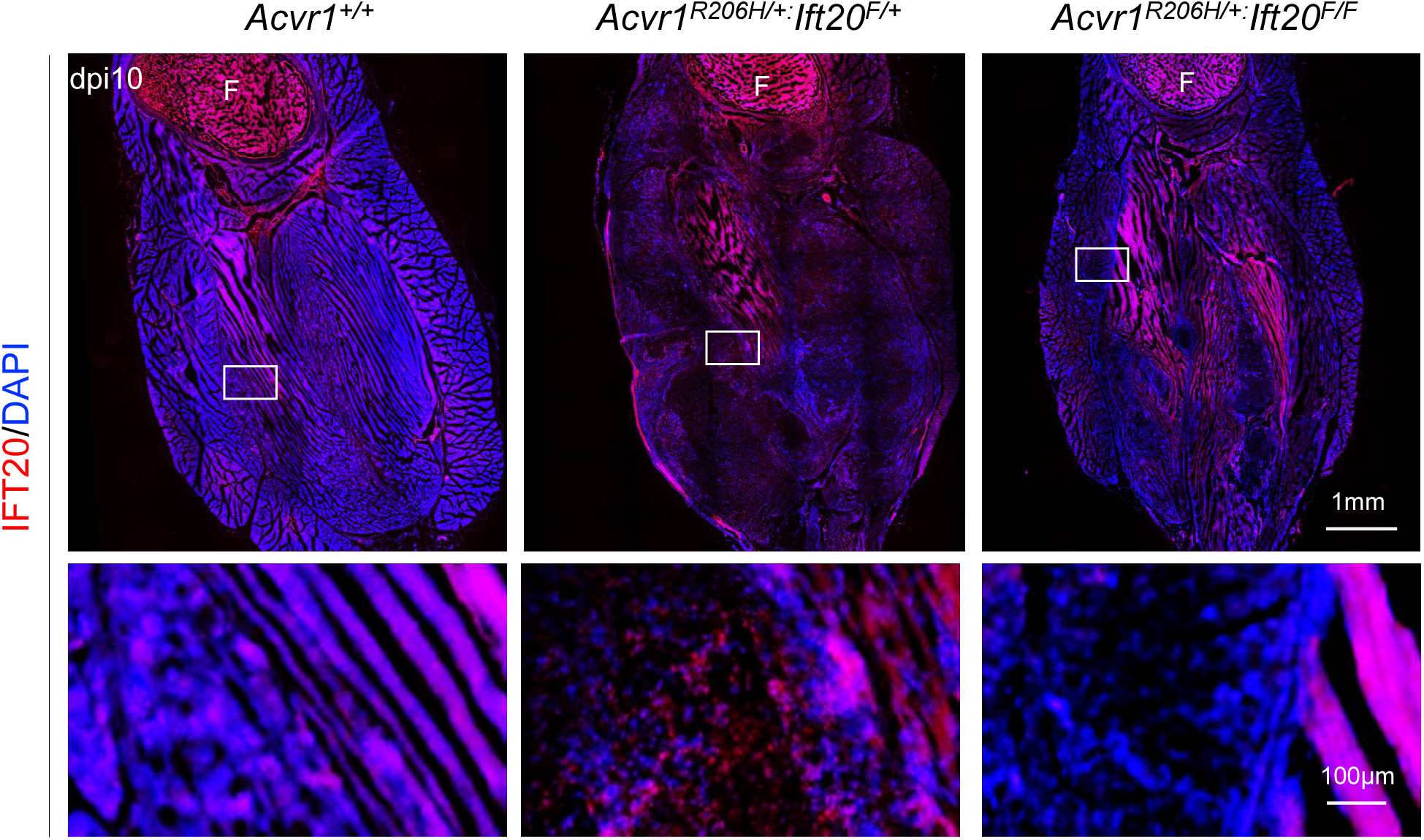
Low magnification images of IFT20. Immunohistochemistry images with anti-IFT20 antibody at dpi10. Lower panels are high-magnification images from the boxed area in the upper panels. F: femur.

**Supplemental Figure 5.**
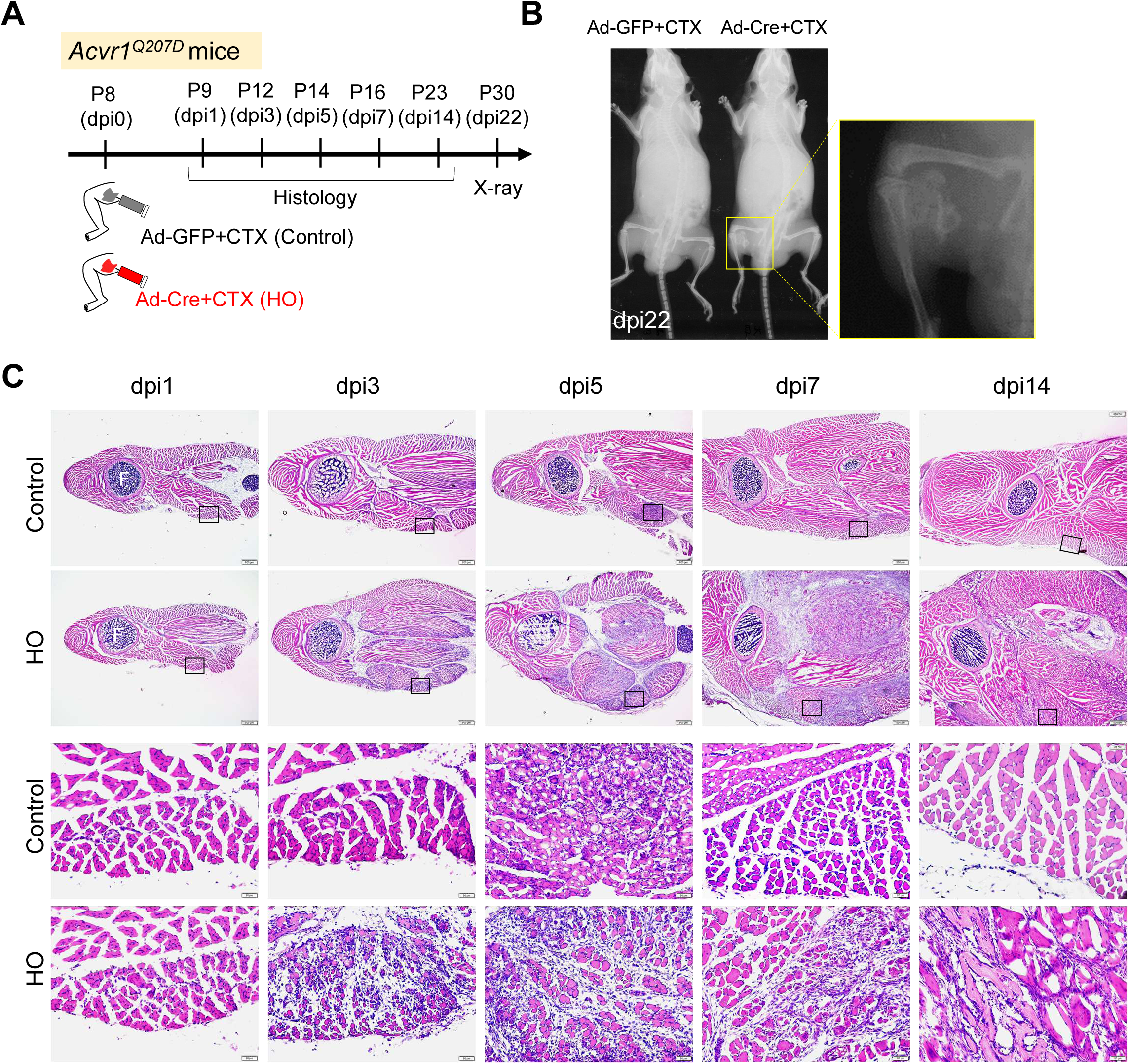
H&E staining in *Acvr1^Q207D^* mice. **(A)** Experimental schedule. **(B)** X-ray images at dpi22. **(C)** Images of H&E staining at dpi1, 3, 5, 7, and 14. Lower panels are high-magnification images from the boxed area (damaged area) in the upper panels. F: femur.

**Supplemental Figure 6.**
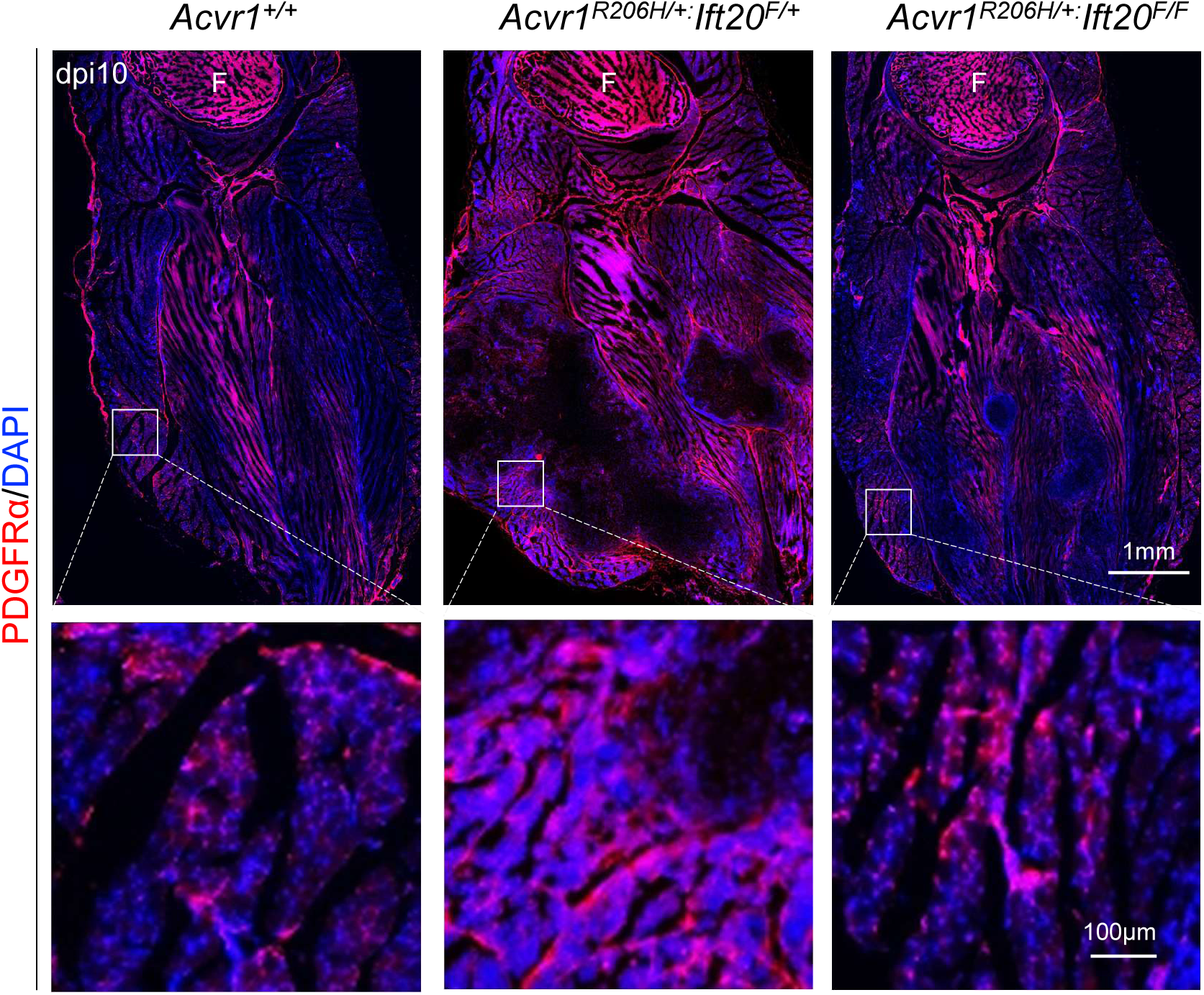
Low-magnification images of FAPs. Immunohistochemistry images with anti-PDGFRα antibody at dpi10. Lower panels are high-magnification images from the boxed area in the upper panels. F: femur.

**Supplemental Figure 7.**
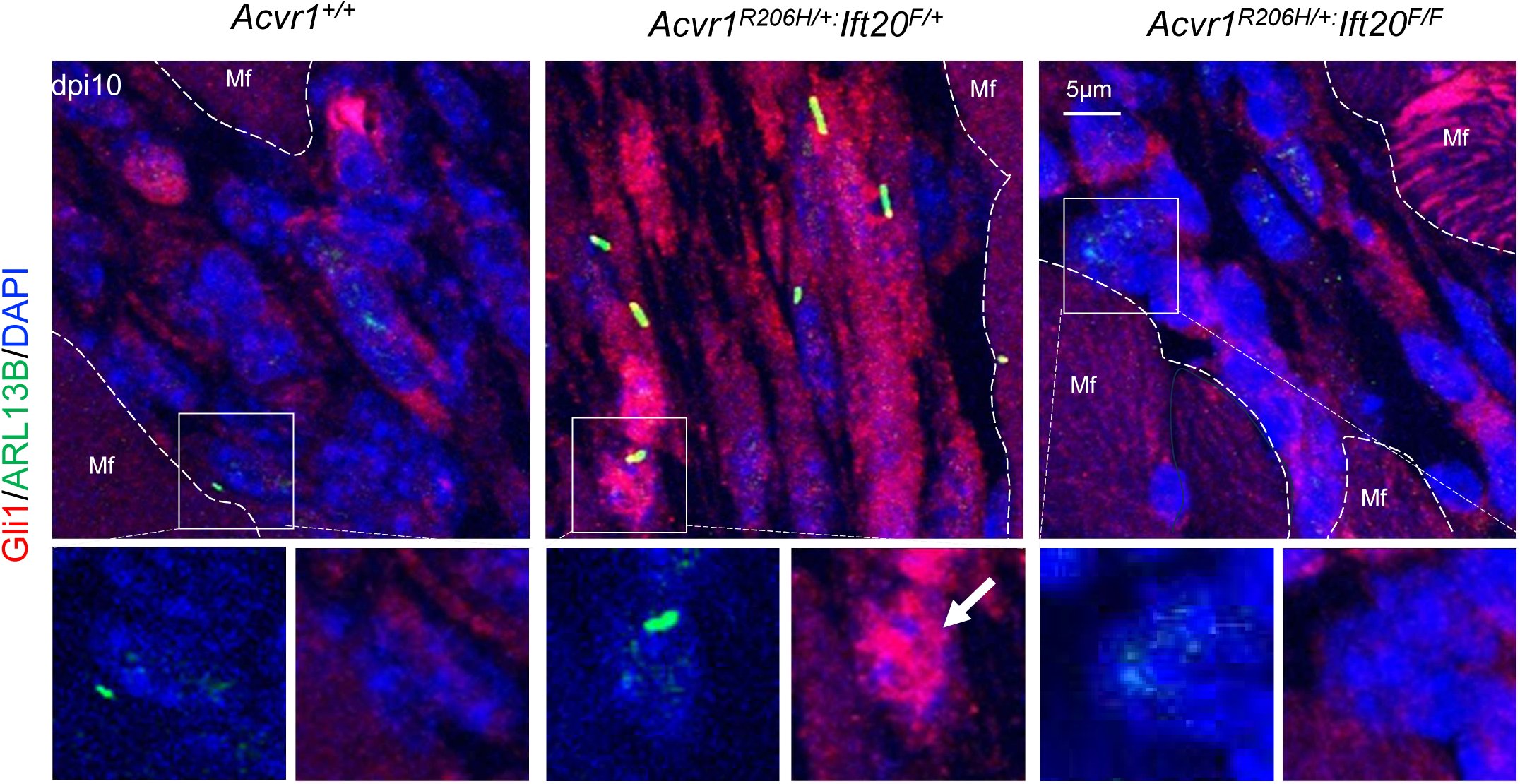
Original images for the Gli1 3D reconstruction. Immunohistochemistry images with anti-GLI1 (red) and anti-ARL13B (green) antibodies at dpi10. Lower panels are high-magnification images from the upper panels. The arrow indicates nuclear localization of GLI1. Mf: muscle fiber.

**Supplemental Table 1.**
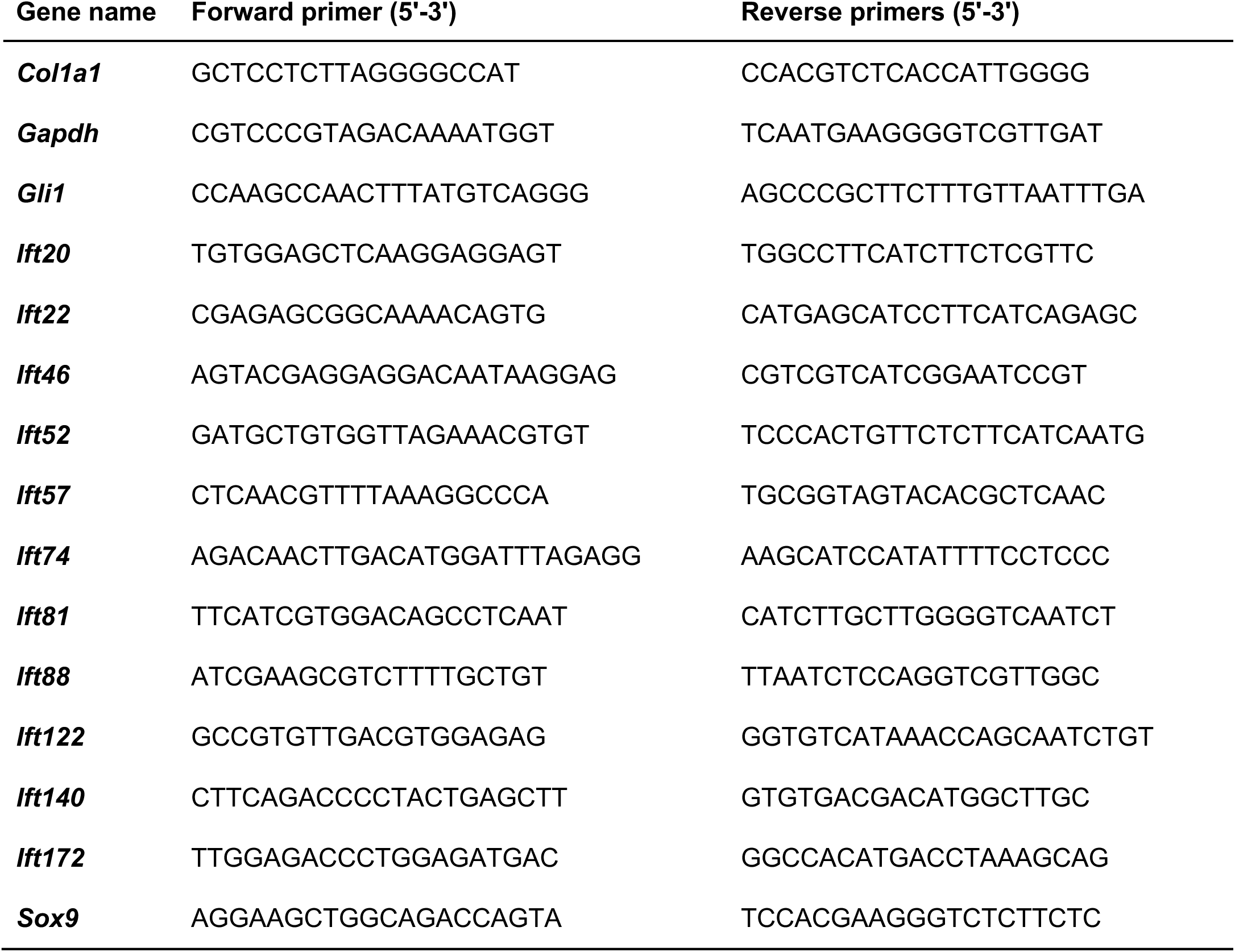
Primer sequence for quantitative RT-PCR.

## References

1. F. S. Kaplan, R. J. Pignolo, E. M. Shore, Heterotopic Ossification: The Keys to the Kingdom. Bone 109, 1–2 (2018).

2. S. Agarwal, M. Sorkin, B. Levi, Heterotopic Ossification and Hypertrophic Scars. Clin Plast Surg 44, 749–755 (2017).

3. F. S. Kaplan et al., Skeletal metamorphosis in fibrodysplasia ossificans progressiva (FOP). Journal of bone and mineral metabolism 26, 521–530 (2008).

4. E. M. Shore et al., A recurrent mutation in the BMP type I receptor ACVR1 causes inherited and sporadic fibrodysplasia ossificans progressiva. Nat Genet 38, 525–527 (2006).

5. K. Shimono et al., Potent inhibition of heterotopic ossification by nuclear retinoic acid receptor-gamma agonists. Nature medicine 17, 454–460 (2011).

6. S. J. Hatsell et al., ACVR1R206H receptor mutation causes fibrodysplasia ossificans progressiva by imparting responsiveness to activin A. Sci Transl Med 7, 303ra137 (2015).

7. J. B. Lees-Shepard et al., Activin-dependent signaling in fibro/adipogenic progenitors causes fibrodysplasia ossificans progressiva. Nat Commun 9, 471 (2018).

8. M. Pacifici, Retinoid roles and action in skeletal development and growth provide the rationale for an ongoing heterotopic ossification prevention trial. Bone 109, 267–275 (2018).

9. R. J. Pignolo, F. S. Kaplan, Clinical staging of Fibrodysplasia Ossificans Progressiva (FOP). Bone 109, 111–114 (2018).

10. Y. S. Yang et al., Suppression of heterotopic ossification in fibrodysplasia ossificans progressiva using AAV gene delivery. Nature communications 13, 6175 (2022).

11. R. J. Pignolo et al., Palovarotene for Fibrodysplasia Ossificans Progressiva (FOP): Results of a Randomized, Placebo-Controlled, Double-Blind Phase 2 Trial. J Bone Miner Res 37, 1891–1902 (2022).

12. R. J. Pignolo et al., Reduction of New Heterotopic Ossification (HO) in the Open-Label, Phase 3 MOVE Trial of Palovarotene for Fibrodysplasia Ossificans Progressiva (FOP). Journal of bone and mineral research : the official journal of the American Society for Bone and Mineral Research 38, 381–394 (2023).

13. R. J. Pignolo et al., Study methodology and insights from the palovarotene clinical development program in fibrodysplasia ossificans progressiva. BMC Med Res Methodol 23, 269 (2023).

14. M. R. Urist, Bone: formation by autoinduction. Science 150, 893–899 (1965).

15. V. S. Salazar, L. W. Gamer, V. Rosen, BMP signalling in skeletal development, disease and repair. Nat Rev Endocrinol 12, 203–221 (2016).

16. K. M. Lyons, V. Rosen, BMPs, TGFbeta, and border security at the interzone. Current topics in developmental biology 133, 153–170 (2019).

17. J. L. Rosenbaum, G. B. Witman, Intraflagellar transport. Nat Rev Mol Cell Biol 3, 813–825 (2002).

18. N. Sharma, N. F. Berbari, B. K. Yoder, Ciliary dysfunction in developmental abnormalities and diseases. Curr Top Dev Biol 85, 371–427 (2008).

19. V. Singla, J. F. Reiter, The primary cilium as the cell’s antenna: signaling at a sensory organelle. Science 313, 629–633 (2006).

20. J. T. Eggenschwiler, K. V. Anderson, Cilia and developmental signaling. Annu Rev Cell Dev Biol 23, 345–373 (2007).

21. J. F. Reiter, M. R. Leroux, Genes and molecular pathways underpinning ciliopathies. Nat Rev Mol Cell Biol 18, 533–547 (2017).

22. B. Basu, M. Brueckner, Fibroblast “cilia growth” factor in the development of left-right asymmetry. Developmental cell 16, 489–490 (2009).

23. S. K. Hong, I. B. Dawid, FGF-dependent left-right asymmetry patterning in zebrafish is mediated by Ier2 and Fibp1. Proceedings of the National Academy of Sciences of the United States of America 106, 2230–2235 (2009).

24. J. M. Neugebauer, J. D. Amack, A. G. Peterson, B. W. Bisgrove, H. J. Yost, FGF signalling during embryo development regulates cilia length in diverse epithelia. Nature 458, 651–654 (2009).

25. M. Kunova Bosakova et al., Regulation of ciliary function by fibroblast growth factor signaling identifies FGFR3-related disorders achondroplasia and thanatophoric dysplasia as ciliopathies. Hum Mol Genet 27, 1093–1105 (2018).

26. L. Martin et al., Constitutively-active FGFR3 disrupts primary cilium length and IFT20 trafficking in various chondrocyte models of achondroplasia. Human molecular genetics 27, 1–13 (2018).

27. G. J. Pazour, J. L. Rosenbaum, Intraflagellar transport and cilia-dependent diseases. Trends Cell Biol 12, 551–555 (2002).

28. T. Fukuda et al., Generation of a mouse with conditionally activated signaling through the BMP receptor, ALK2. Genesis 44, 159–167 (2006).

29. Y. Komatsu, V. Kaartinen, Y. Mishina, Cell cycle arrest in node cells governs ciliogenesis at the node to break left-right symmetry. Development 138, 3915–3920 (2011).

30. Y. Komatsu, Y. Mishina, Establishment of left-right asymmetry in vertebrate development: the node in mouse embryos. Cell Mol Life Sci 70, 4659–4666 (2013).

31. T. J. van Dam, G. Wheway, G. G. Slaats, M. A. Huynen, R. H. Giles, The SYSCILIA gold standard (SCGSv1) of known ciliary components and its applications within a systems biology consortium. Cilia 2, 7 (2013).

32. J. B. Regard et al., Activation of Hedgehog signaling by loss of GNAS causes heterotopic ossification. Nature medicine 19, 1505–1512 (2013).

33. Q. Cong et al., A self-amplifying loop of YAP and SHH drives formation and expansion of heterotopic ossification. Sci Transl Med 13 (2021).

34. K. G. Kozminski, K. A. Johnson, P. Forscher, J. L. Rosenbaum, A motility in the eukaryotic flagellum unrelated to flagellar beating. Proceedings of the National Academy of Sciences of the United States of America 90, 5519–5523 (1993).

35. L. J. Jonk, S. Itoh, C. H. Heldin, P. ten Dijke, W. Kruijer, Identification and functional characterization of a Smad binding element (SBE) in the JunB promoter that acts as a transforming growth factor-beta, activin, and bone morphogenetic protein-inducible enhancer. The Journal of biological chemistry 273, 21145–21152 (1998).

36. K. Kusanagi et al., Characterization of a bone morphogenetic protein-responsive Smad-binding element. Molecular biology of the cell 11, 555–565 (2000).

37. K. C. Corbit et al., Vertebrate Smoothened functions at the primary cilium. Nature 437, 1018–1021 (2005).

38. K. C. Corbit et al., Kif3a constrains beta-catenin-dependent Wnt signalling through dual ciliary and non-ciliary mechanisms. Nat Cell Biol 10, 70–76 (2008).

39. P. J. Ocbina, M. Tuson, K. V. Anderson, Primary cilia are not required for normal canonical Wnt signaling in the mouse embryo. PLoS One 4, e6839 (2009).

40. D. Huangfu, K. V. Anderson, Cilia and Hedgehog responsiveness in the mouse. Proc Natl Acad Sci U S A 102, 11325–11330 (2005).

41. S. C. Goetz, K. V. Anderson, The primary cilium: a signalling centre during vertebrate development. Nat Rev Genet 11, 331–344 (2010).

42. A. J. Firestone et al., Small-molecule inhibitors of the AAA+ ATPase motor cytoplasmic dynein. Nature 484, 125–129 (2012).

43. D. H. Roossien, K. E. Miller, G. Gallo, Ciliobrevins as tools for studying dynein motor function. Front Cell Neurosci 9, 252 (2015).

44. L. Sun et al., Oxidative phosphorylation is a pivotal therapeutic target of fibrodysplasia ossificans progressiva. Life Sci Alliance 7 (2024).

45. M. N. Wosczyna, A. A. Biswas, C. A. Cogswell, D. J. Goldhamer, Multipotent progenitors resident in the skeletal muscle interstitium exhibit robust BMP-dependent osteogenic activity and mediate heterotopic ossification. Journal of bone and mineral research : the official journal of the American Society for Bone and Mineral Research 27, 1004–1017 (2012).

46. A. W. Joe et al., Muscle injury activates resident fibro/adipogenic progenitors that facilitate myogenesis. Nature cell biology 12, 153–163 (2010).

47. M. N. Wosczyna et al., Mesenchymal Stromal Cells Are Required for Regeneration and Homeostatic Maintenance of Skeletal Muscle. Cell reports 27, 2029–2035 e2025 (2019).

48. M. M. Murphy, J. A. Lawson, S. J. Mathew, D. A. Hutcheson, G. Kardon, Satellite cells, connective tissue fibroblasts and their interactions are crucial for muscle regeneration. Development 138, 3625–3637 (2011).

49. K. He et al., Primary cilia mediate skeletogenic BMP and Hedgehog signaling in heterotopic ossification. Sci Transl Med 16, eabn3486 (2024).

50. B. Lai et al., MEST inhibits ciliary sphingomyelin synthesis to promote tendon stem/progenitor cell osteochondrogenesis in traumatic heterotopic ossification. Genes & Diseases 10.1016/j.gendis.2026.102166, 102166 (2026).

51. C. A. Clement et al., TGF-beta signaling is associated with endocytosis at the pocket region of the primary cilium. Cell reports 3, 1806–1814 (2013).

52. P. Mill, S. T. Christensen, L. B. Pedersen, Primary cilia as dynamic and diverse signalling hubs in development and disease. Nature reviews. Genetics 24, 421–441 (2023).

53. D. Zhang et al., ALK2 functions as a BMP type I receptor and induces Indian hedgehog in chondrocytes during skeletal development. Journal of bone and mineral research : the official journal of the American Society for Bone and Mineral Research 18, 1593–1604 (2003).

54. D. Kopinke, E. C. Roberson, J. F. Reiter, Ciliary Hedgehog Signaling Restricts Injury-Induced Adipogenesis. Cell 170, 340–351 e312 (2017).

55. T. Molina, P. Fabre, N. A. Dumont, Fibro-adipogenic progenitors in skeletal muscle homeostasis, regeneration and diseases. Open Biol 11, 210110 (2021).

56. B. C. Collins, G. Kardon, It takes all kinds: heterogeneity among satellite cells and fibro-adipogenic progenitors during skeletal muscle regeneration. Development 148 (2021).

57. A. Stanley et al., Dynamics of skeletal muscle-resident stem cells during myogenesis in fibrodysplasia ossificans progressiva. NPJ Regen Med 7, 5 (2022).

58. F. Long, X. M. Zhang, S. Karp, Y. Yang, A. P. McMahon, Genetic manipulation of hedgehog signaling in the endochondral skeleton reveals a direct role in the regulation of chondrocyte proliferation. Development 128, 5099–5108 (2001).

59. J. A. Jonassen, J. San Agustin, J. A. Follit, G. J. Pazour, Deletion of IFT20 in the mouse kidney causes misorientation of the mitotic spindle and cystic kidney disease. The Journal of cell biology 183, 377–384 (2008).

60. H. Yamaguchi et al., Temporospatial regulation of intraflagellar transport is required for the endochondral ossification in mice. Dev Biol 482, 91–100 (2022).

61. K. Noda, Y. Mishina, Y. Komatsu, Constitutively active mutation of ACVR1 in oral epithelium causes submucous cleft palate in mice. Developmental biology 10.1016/j.ydbio.2015.06.014 (2015).

62. S. S. V. Vasquez, J. van Dam, G. Wheway, An updated SYSCILIA gold standard (SCGSv2) of known ciliary genes, revealing the vast progress that has been made in the cilia research field. Mol Biol Cell 32, br13 (2021).

63. R. Patro, G. Duggal, M. I. Love, R. A. Irizarry, C. Kingsford, Salmon provides fast and bias-aware quantification of transcript expression. Nat Methods 14, 417–419 (2017).

64. M. I. Love, W. Huber, S. Anders, Moderated estimation of fold change and dispersion for RNA-seq data with DESeq2. Genome Biol 15, 550 (2014).

